# Abolishing respiratory complex I decreases in vivo growth of high grade serous ovarian cancer cells and sensitizes to anti-angiogenic therapy

**DOI:** 10.64898/2026.02.28.708681

**Authors:** Ivana Kurelac, Beatrice Cavina, Francesca Nanetti, Simona Corrà, Manuela Sollazzo, Camelia Alexandra Coada, Marco Grillini, Laura Scalambra, Eleonora Lama, Eleonora Angi, Sonia Minuzzo, Luisa Iommarini, Stefano Indraccolo, Anna Maria Porcelli, Giuseppe Gasparre

**Affiliations:** Department of Medical and Surgical Sciences (DIMEC), University of Bologna, Bologna, Italy; IRCCS Azienda Ospedaliero-Universitaria Di Bologna, Bologna, Italy; Department of Pharmacy and Biotechnology (FABIT), University of Bologna, Bologna, Italy; Division of Physiology, Department of Morpho-Functional Sciences, University of Medicine and Pharmacy “Iuliu Hațieganu”, Cluj-Napoca, Romania; Pathology Unit, IRCCS Azienda Ospedaliero-Universitaria di Bologna, Bologna, Italy; Center for Applied Biomedical Research (CRBA), University of Bologna, Bologna, Italy; Department of Surgery, Oncology and Gastroenterology, University of Padua, Padua, Italy; Basic and Translational Oncology Unit, Veneto Institute of Oncology IOV-IRCCS, Padua, Italy; Centro Studi e Ricerca Sulle Neoplasie Ginecologiche, University of Bologna, Bologna, Italy

**Keywords:** Mitochondrial Complex I, high grade serous tubo-ovarian cancer (HGSOC), bevacizumab, hypoxia inducible factor-1 (HIF-1)

## Abstract

Targeting mitochondrial Complex I (CI) is a currently emerging anti-cancer strategy, with several enzyme inhibitors entering clinical trials. Among others, aggressive high-grade serous tubo-ovarian cancer (HGSOC) may particularly benefit from this therapeutic approach due to the scarce response to first- and second-line treatments, with consequent high mortality, such as the anti-angiogenic bevacizumab. We here show that CI represents a vulnerability in HGSOC, which can be exploited for therapeutic intervention. Indeed, ablating CI function in OV-90 HGSOC cells led to significant *in vivo* tumor growth decrease, smaller masses, and lower KI-67 proliferative index. This was confirmed in a switch-off system in which CI deprivation was induced during tumor progression to mimic pharmacologic treatment, suggesting this result can be achieved in growing neoplasms. We also show that abolishing CI in HGSOC cells leads to failure in stabilizing the hypoxia inducible factor-1a and to respond to hypoxia through the transcriptional activation of its target genes, ultimately lowering vascular endothelial growth factor (VEGF) and generating an immature intratumor vascular system accompanied by a decreased blood flow. Last, we demonstrate that targeting CI sets the biological basis for increased sensitivity to anti-angiogenics, as CI-deprived tumors displayed growth arrest when bevacizumab was administered, unlike their CI-competent counterpart. Our findings point to CI inhibition as a booster for anti-VEGF therapies and pave the way for combined protocols in treatment of HGSOC.

## Introduction

Mitochondrial respiratory Complex I (CI) is a multi-subunit enzyme whose crucial role in many metabolism-related cellular processes, among which ATP production, aspartate biosynthesis and response to hypoxia^1–3^, has placed it in the spotlight in oncology as its dysfunction represents a vulnerability for cancer cells. Several CI inhibitors have indeed been developed in the search for potential anticancer therapies, some of which have been clinically tried against solid or non-solid tumors (i.e. IACS, metformin^4,5^). Only recently, targeting CI has been undertaken also in ovarian cancer (OC) both in preclinical and clinical contexts^6,7^. This is a cogent issue particularly for the high-grade serous tubo-ovarian (HGSOC) histotype, which presents high recurrence rate, chemoresistance onset, and, ultimately, high disease-associated mortality^8,9^. While inconclusive results have been obtained in trials, likely due to the use of low specificity CI inhibitors such as metformin, preclinical studies show potential for CI targeting efficacy, provided adaptive mechanisms are prevented and subcategories of vulnerable OCs are identified by means of specific metabolic or molecular features. We previously exploited CI knockouts, both stable and inducible, to delve into the cellular responses that solid cancers implement following enzyme dysfunction and the consequent energetic impairment, to unveil vulnerabilities that may be targeted in combinatorial strategies to increase efficacy, such as macrophage inhibition^10^. One such response we demonstrated to occur horizontally in osteosarcoma, colorectal, and thyroid cancer is the inability of CI-deprived cells to properly sense hypoxia and trigger stabilization of hypoxia inducible factor-1α (HIF1a), with subsequent lower expression of cell autonomous vascular endothelial growth factor (VEGF) and development of an alternative intratumoral vasculature system characterized by lack of pericytes^10^. We here apply the same approach to OC, taking advantage of a highly aggressive HGSOC-derived cell line with the ability to form large tumors in preclinical models, and show that CI abolishment holds promise to help contain the disease also in ovarian neoplasia. Moreover, since anti-angiogenic bevacizumab (BEV) is administered in HGSOC as part of maintenance therapy protocols with limited increase in both progression-free and overall survival, we show that CI dysfunction synergizes with anti-VEGF therapy in mice to halt disease progression, which may be linked to the defective HIF-1/VEGF phenotype these tumors display. We believe our results pave the way for the combined use of CI inhibitors with anti-VEGF drugs to boost efficacy of therapeutic protocols in HGSOC care.

## Results

We previously demonstrated that knock-out (KO) of CI subunit NDUFS3 abolishes enzymatic activity^10–12^ and curbs tumorigenic potential *in vivo* of colorectal and osteosarcoma cells^10^. We here translate those findings into OC. *NDUFS3* KO was performed by CRISPR/Cas-9, as previously described^12^, in HGSOC-derived OV-90 cells, which present the advantage of growing well in NOD/SCID mice, unlike other commonly used OC cell lines, allowing us to test their tumorigenic potential *in vivo*. Hence, CI-competent and *NDUFS3* KO OV-90 cells (OV-90^+/+^ and OV-90^−/−^, respectively) were injected subcutaneously in NOD/SCID mice and tumor growth was monitored over time. At day 29 after injection, the first mouse of our cohort reached the experimental end point, and the experiment was terminated. The average tumor mass volume of the animals injected with OV-90^−/−^ was lower than that of animals carrying OV-90^+/+^ masses (p = 0.0096, one-tailed t-test, Fig. 1A). This difference was significant also on day 27 post-injection (p = 0.0362, one-tailed t-test, Fig. 1A). Excised tumor volumes as well were concordantly lower in the CI-KO group (Fig. 1B), clearly suggesting that CI-ablated OC cells display a lower tumorigenic potential and a slower growth *in vivo*, similarly to other cancer types^10^. To ascertain that xenografts had a phenotype that reflected the genetic *NDUFS3* KO, we performed immunohistochemistry (IHC) with anti-NDUFS3 antibody, revealing a homogeneously negative staining in OV-90^−/−^ masses compared with positive staining in their CI competent counterpart (Fig. 1C). A negative staining for CI subunit NDUFS4 in NDUFS3-depleted xenografts exclusively (Fig. 1C) indicated that CI assembly had indeed been impaired^13^. These data were further confirmed by western blot analysis using a human-specific NDUFS3 antibody, which showed the absence of the subunit in the explanted xenografts (Fig. 1D). The KI-67 mitotic index of KO tumors was significantly lower compared to CI-competent xenografts (Fig. 1E), suggesting that the growth slowdown observed *in vivo* was due to decreased proliferative capacity of OV-90^−/−^ cells. To unequivocally demonstrate that the growth impairment was due to the lack of NDUFS3 and not due to a nonspecific, acquired fragility of the KO cells generated, we stably re-expressed the subunit in OV-90^−/−^ cells (Fig. 1F) and injected the latter, from here onwards named OV-90^−/−S3^, in NOD/SCID mice along with their mock controls (OV-90^−/−M^). As expected, OV-90^−/−S3^ tumors grew faster than their mock counterpart, and their excised masses were concordantly larger (Fig. 1G-H), allowing us to conclude that the lack of NDUFS3 and consequent abolishment of CI could be appointed for the dampening of OV-90 tumor growth.

**Figure 1.**
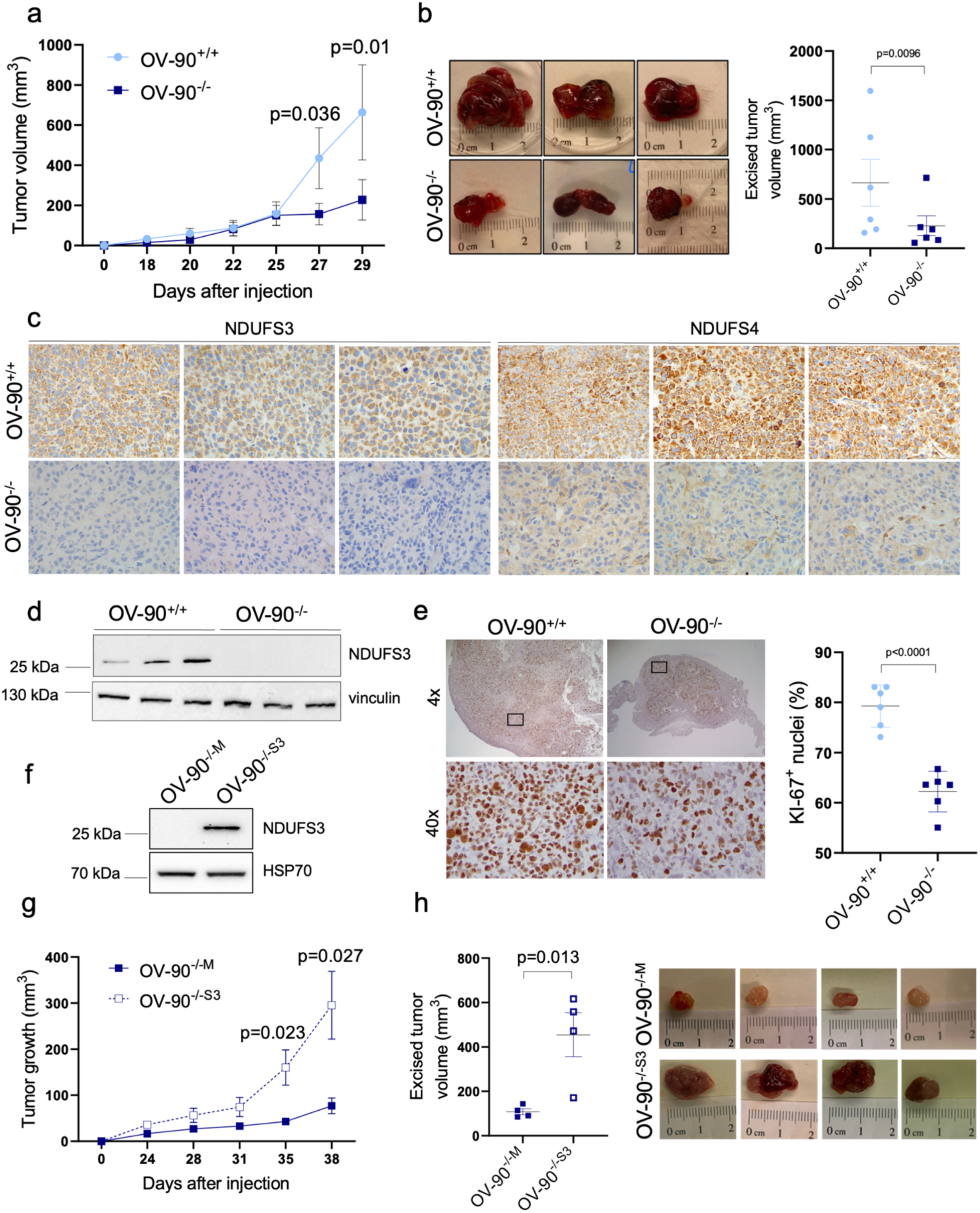
NDUFS3 knock-out causes reduction of OC tumorigenic potential. **(a)** Growth curves of wild type (OV-90^+/+^) and NDUFS3 KO (OV-90^−/−^) OC xenografts in NOD/SCID mice (n=6 per group). One-tailed t-test was used to compare averages between the groups for each day of measurement. F-test was significant for day 27 and day 29. Thus, the data were transformed (y’=logy) prior to t-test calculation. **(b)** Excised tumor volumes of OV-90^+/+^and OV-90^−/−^ xenografts. One-tailed t-test was used to compare averages. Grubbs test identified one outlier which was excluded from statistical testing. F-test identified significant differences. Thus, data were transformed (y’=logy) prior to t-test calculation. Representative tumors are shown. **(c)** IHC for NDUFS3 and NDUFS4 CI subunits. Representative images are shown. **(d)** NDUFS3 western blot analysis in OV-90 xenografts (n=3 per group). Vinculin was used as loading control. **(e)** Representative images of KI-67 IHC staining and quantification of cells displaying KI-67^+^ nuclei in OV-90 tumors. Representative images are shown. Black squares indicate the insets of the areas shown at higher magnification. **(f)** NDUFS3 western blot analysis in OV-90^−/−^ cells stably transduced with empty (OV-90^−/−M^) and NDUFS3 carrying vector (OV-90^−/−S3^). HSP70 was used as loading control. **(g)** Growth curves of OV-90^−/−M^ and OV-90^−/−S3^ xenografts in NOD/SCID mice (n=4 per group). **(h)** Excised tumor volumes of OV-90^−/−M^ and OV-90^−/−S3^ xenografts. Representative tumors are shown. For all panels, data in graphs are represented as mean ± SEM and p-value is indicated.

Overall, we were able to recapitulate in OC setting the same anti-tumorigenic phenotype induced upon *NDUFS3* KO and CI ablation previously observed in other neoplastic contexts, which prompted us to test whether it was feasible to trigger a growth slowdown in progressing masses, through the use of a negative inducible system where re-expressed NDUFS3 in OV-90^−/−^ cells may be switched off by doxycycline (DOX) administration. Indeed, the *NDUFS3* transgene carried by the plasmidic construct transduced in OV-90 KO cells was under negative control by DOX, similar to what we previously reported^10^. The system was validated in terms of appropriate NDUFS3 re-expression and switch-off when DOX was administered *in vitro* at doses that do not affect general mitochondrial protein synthesis (Fig. 2A). We also monitored the disappearance of CI over time to gauge the turnover of the complex as a whole and revealed that 20 days of treatment were required for complete loss of CI activity (Fig. 2B). Next, the cells were injected in NOD/SCID mice and animals carrying OV-90^−/−S3^ tumors displayed shorter survival compared to OV-90^−/−M^ (Fig. 2C). At the same time, we attempted to evaluate whether the tumorigenic potential of OV-90 cells may be blunted when CI assembly is impaired during cancer progression *in vivo*. As expected, OV-90^−/−S3^ tumors that had been randomized to receive DOX indeed reached the experimental end point in terms of mass volume more slowly, resulting in prolonged animal survival upon Kaplan-Meier analysis (Fig. 2D). IHC staining for NDUFS3 in explanted xenografts showed that protein expression had been appropriately recovered in OV-90^−/−S3^ tumors compared to mock, as well as completely abolished in DOX-treated animals (Fig. 2E). Concordantly, the expression levels of NDUFS4, as a marker of CI assembly, were recuperated in OV-90^−/−S3^ tumors compared to mock and significantly lower in DOX-treated masses (Fig. 2E) and CI activity was markedly evident only in OV-90^−/−S3^ tumors (Fig. 2F). Last, to infer on the more or less indolent behavior, we measured the mitotic index of the three groups and revealed that KI-67 was significantly higher in OV-90^−/−S3^ tumors compared to both mock and DOX-treated ones (Fig. 2G), confirming that switching off *NDUFS3* expression during cancer growth curbs the replicating ability of cancer cells, and overall demonstrating the anti-tumorigenic effect of impairing CI during OC progression.

**Figure 2.**
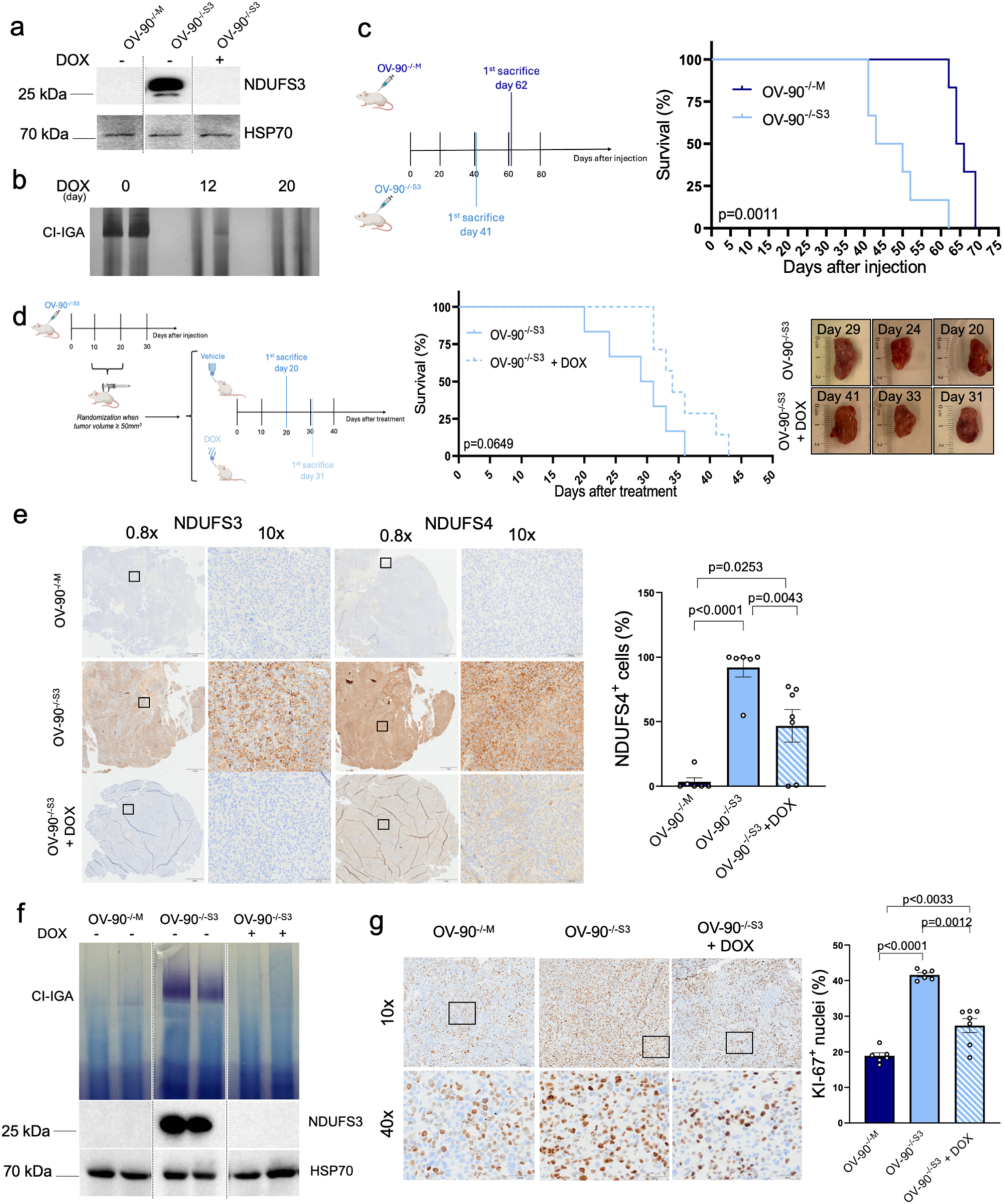
Targeting CI slows down OC progression. **(a)** NDUFS3 western blot in OV-90^−/−^ cells stably transduced with empty (OV-90^−/−M^) or NDUFS3 carrying vector (OV-90^−/−S3^) and cultured with or without DOX (100 ng/mL) for 12 days. HSP70 was used as loading control. **(b)** CI In-Gel Activity (CI-IGA) in OV-90^−/−S3^ cells cultured with or without DOX (100 ng/mL) at indicated time points. A representative gel of three independent experiments is shown. **(c)** Graphical sketch of the experimental setting and Kaplan-Meier curve of NOD/SCID mice injected with OV-90^−/−M^ or OV-90^−/−S3^ cells (n=6 per group). Survival end-point: xenografts reaching 500 mm^3^. **(d)** Graphical sketch of the experimental setting and Kaplan-Meier curve of NOD/SCID mice injected with OV-90^−/−S3^ cells and randomized when reaching 50 mm^3^ in groups treated with (n=7) or without (n=6) DOX (1 mg/mL in drinking water). Representative tumors are shown. Survival end-point: xenografts reaching 500 mm^3^. **(e)** IHC for NDUFS3 and NDUFS4 CI subunits. Representative images are shown, together with the quantification of NDUFS4 positivity. Black squares indicate the insets of the areas shown at higher magnification. Grubbs test identified outliers which were excluded from statistical testing. Shapiro-Wilk test revealed non-normal distribution of OV-90^−/−M^ and OV-90^−/−S3^ data. Thus, Mann-Whitney test was used to compare the groups. **(f)** CI-IGA and NDUFS3 western blot in OV-90^−/−M^ and OV-90^−/−S3^ masses derived from NOD/SCID mice treated with or without DOX (1 mg/mL in drinking water). A representative gel of n=2 xenografts per group is shown. HSP70 was used as loading control. **(g)** Quantification of cells displaying KI-67 positive nuclei in OV-90 tumors. Shapiro-Wilk test revealed non-normal distribution of OV-90^−/−S3^+DOX data. Thus, Mann-Whitney test was used to compare the groups. Representative images of KI-67 IHC staining are shown with black squares indicating the insets of the areas shown at higher magnification. For Kaplan-Meier curves, Log-rank (Mantel-Cox) test was performed to compare the groups and p-value is indicated. For all other panels, data in graphs are represented as mean ± SEM and p-value is indicated.

We next delved into the molecular phenotype of pseudo-normoxia we previously observed in other cancers to occur by means of chronic HIF1a destabilization, following CI disruption. Indeed, western blotting for HIF1a in explanted tumors showed a marked decrease or lack of this transcription factor in KO masses compared to WT (Fig. 3A), regardless of their size, suggesting a general inability to trigger a proper HIF1-driven response when CI is ablated. Concordantly, the expression of a panel of genes that are positively regulated by HIF1 at the transcriptional level (glucose transporter - *SLC2A1*, lactate dehydrogenase A – *LDHA*, Macrophage Inhibitory Factor - *MIF* and vascular endothelial growth factor A – *VEGFA*) was decreased in CI-KO tumors compared to WT (Fig. 3B). These outcomes prompted us to explore the vasculature characteristics, as we have previously shown both in colorectal cancer and osteosarcoma that CI disruption leads to a slower growth that must rely on the host stroma for nutrient supply, which is supported by the scarcity of mature vessels permeating the tumor tissue^10^. Similarly to what we had observed in other cancers, OV-90 tumors lacking CI showed a higher abundance of stromal components (Fig. 3C). Moreover, compared to their WT counterparts, the number of vessels (endomucin^+^) was higher in OV-90^−/−^ masses, but they were characterized by lack of pericytes, as gauged by co-staining of endomucin with Smooth Muscle Actin (SMA), which are part of mature vascular structures exclusively (Fig. 3D). In agreement with these findings, intratumoral blood flow was decreased in CI-KO masses (Fig. 3E) despite their higher vessel number. Interestingly, at variance with our previous observations in other solid neoplasms, even though the KO masses presented with significantly higher number of macrophages compared to their WT counterparts (Fig. 3F) and a significant decrease in *MIF* expression (Fig. 3B), the absolute number of tumor infiltrating macrophages was very low, with scant and spot-like appearances of few myeloid cells. This is coherent, however, with the knowledge that most aggressive HGSOC subtypes are immunologically cold^14^.

**Figure 3.**
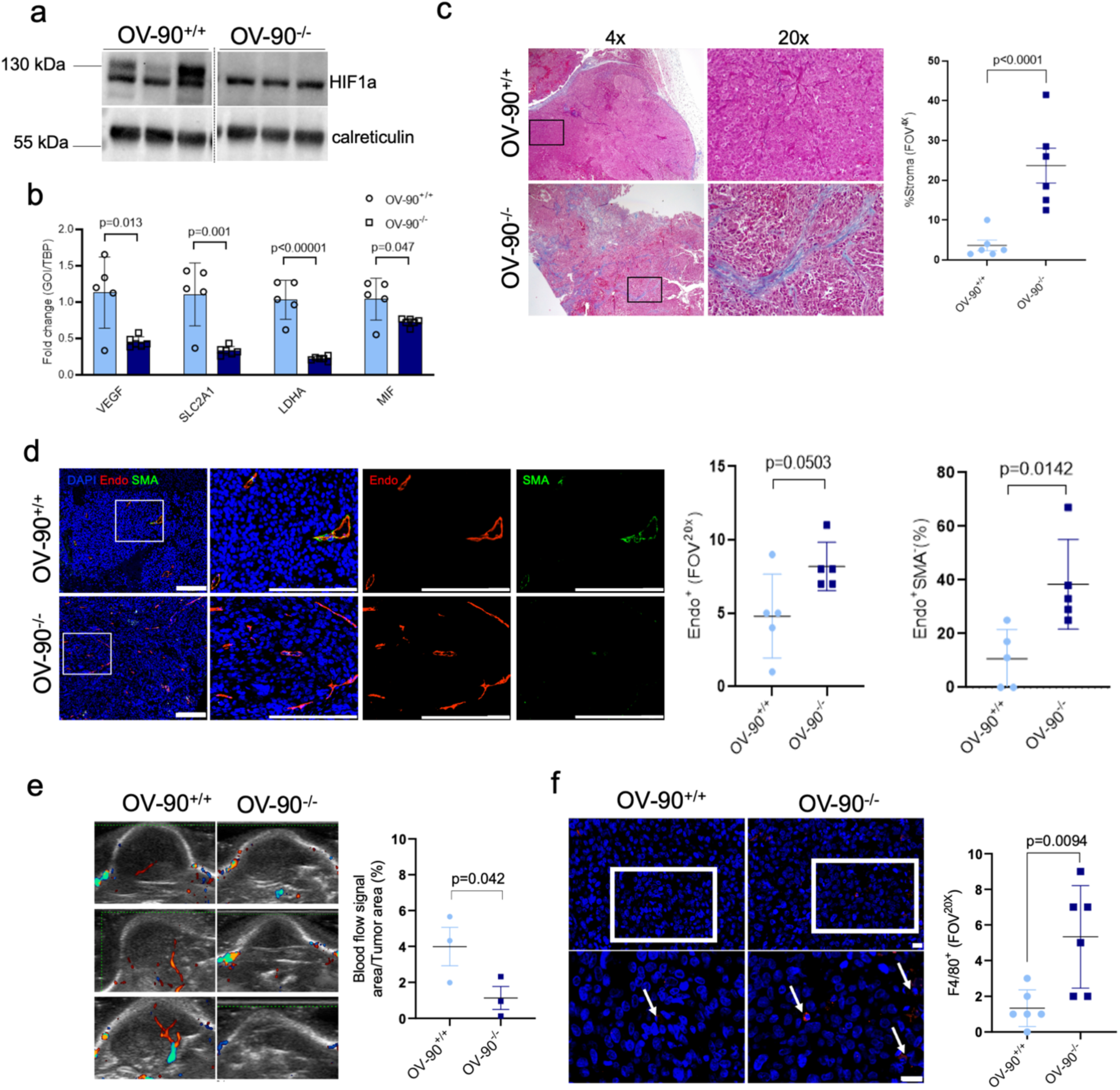
CI deficient OC masses display HIF1 inactivation associated with immature vasculature and decreased intratumoral blood flow. **(a)** HIF1a western blot analysis in representative OV-90^+/+^ and OV90^−/−^ xenografts (n=3 per group). Calreticulin was used as loading control. **(b)** Gene expression of HIF1 targets evaluated by qRT-PCR in OV-90^+/+^(n=5) and OV-90^−/−^(n=6) xenografts. Data were log transformed prior to statistical analysis. **(c)** Masson’s trichrome staining of OV-90^+/+^ and OV90^−/−^ xenografts, with respective quantification of the stroma (collagen areas stained in blue). Representative images are shown with black squares indicating the insets of the areas shown at higher magnification. **(d)** Immunofluorescent staining of endomucin (Endo) and Smooth Muscle Actin (SMA) in OV-90^+/+^ and OV-90^−/−^ xenografts, with nuclei stained with DAPI, scale bar 200 µm. White squares indicate the insets of the areas shown at higher magnification. Representative images are shown, together with quantification of total (Endo^+^) and pericyte negative (Endo^+^SMA^−^) vessels. **(e)** Ultrasound images of OV-90^+/+^ and OV-90^−/−^ xenografts. Incoming (red) and outcoming (blue) blood flows are shown. The graph indicates the percentage of the blood flow areas detected within each tumor (n=3 per group). One-tailed t-test was used to compare averages between the groups. **(f)** Immunofluorescent staining of F4/80^+^ macrophages (red) in OV-90^+/+^ and OV-90^−/−^ xenografts. White squares indicate the insets of the areas shown at higher magnification. The arrows indicate F4/80^+^ cells. Nuclei are stained with DAPI (blue), scale bar = 20 µm. For all panels, data in graphs are represented as mean ± SEM and p-value is indicated.

Taken together, these data prompted us to hypothesize that lack of HIF1a stabilization and the inability to respond to hypoxia that occurs in CI-ablated OCs may render cancer cells more dependent on VEGF-driven neoangiogenesis, in a condition where both cell and non-cell autonomous VEGF is scarcely produced, ultimately rendering CI-KO tumors more sensitive to BEV. This hypothesis was also fostered by our recent data on the increased sensitivity to BEV of OC patient-derived xenografts with dysfunctional oxidative phosphorylation^15^. We therefore challenged OV-90 xenografts, both CI-KO and WT with this anti-angiogenic drug and revealed that KO tumors were significantly more sensitive to VEGF function abolishment than their CI-competent counterparts (Fig. 4A-G). In particular, whereas no treatment effect was observed during early stages on NDUFS3 WT tumors (Fig. 4A), the growth of CI-KO xenografts was significantly affected already at day 7 (*p* = 0.007; Fig. 4B), underlining their evident higher sensitivity. In fact, despite an increase in survival of the OV-90^+/+^ group receiving BEV compared to untreated (*p* = 0.02), all mice carrying CI-competent tumors reached the 500 mm^3^ end point by day 40 (Fig. 4C). On the other hand, strikingly, compared to controls, BEV treated mice carrying OV-90^−/−^ tumors were all still alive, with little or no sign of disease progression and distress up to the 50^th^ day of experiment, i.e. the ethically-defined end timepoint (*p* = 0.0008, Fig. 4D). Concordantly, the median excised volume of the OV-90^−/−^ tumors treated with BEV was 192 mm^3^ (Fig. 4E). Post explant analyses showed that the HIF1a destabilization was maintained in KO versus CI-competent tumors, and unchanged by BEV treatment (Fig. 4F). As expected, BEV significantly decreased the overall number of vessels (endomucin^+^) in CI-competent and KO tumors alike (Fig. 4G). Interestingly, however, while in CI-competent masses the proportion of mature (SMA^+^) versus immature (SMA^−^) vessels did not change, in CI-KO grafts the reduction in vessel number upon treatment was mainly attributable to the decrease of the vessels lacking pericyte coating (Fig. 4G), strongly indicating that BEV had a massive impact in halting the formation of novel, initially immature vessels in tumors that rely predominantly on this type of vasculature for growth and survival. Overall, by allowing persistence exclusively of vessels already formed before the beginning of treatment, BEV efficiently impeded disease progression when CI was lacking and the HIF1/VEGFA axis impaired.

**Figure 4.**
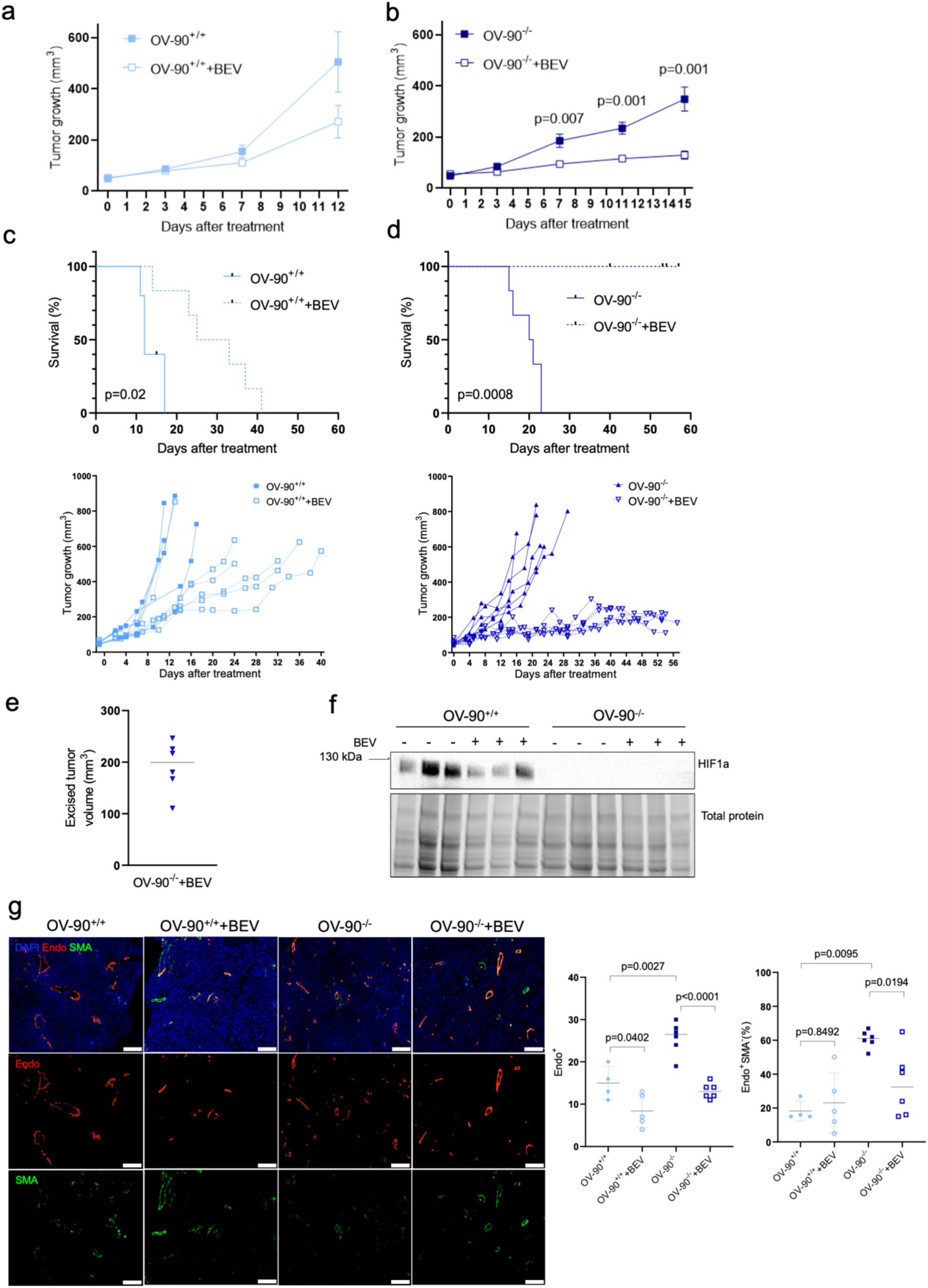
Targeting CI sensitizes OC to anti-angiogenic therapy. **(a)** Growth curves of NDUFS3 wild type (OV-90^+/+^) xenografts in NOD/SCID mice treated with (+BEV, n=4) or without (n=5) BEV. Data are represented as mean ± SEM. **(b)** Growth curves of OV-90^−/−^ xenografts in NOD/SCID mice treated with (+BEV) or without BEV (n=6 per group). Data are represented as mean ± SEM. **(c)** Kaplan-Meier curve of NOD/SCID mice injected with OV-90^+/+^cells, randomized when reaching 50 mm^3^ in groups and treated bi-weekly via intraperitoneal injection with (+BEV, n=5) or without (n=6) BEV (100 µg/dose) and xenograft growth curves of single NOD/SCID mice treated with (+BEV, n=4) or without (n=5) BEV. Survival end-point: xenografts reaching 500 mm^3^. **(d)** Kaplan-Meier curve and xenograft growth curves of NOD/SCID mice injected with OV-90^−/−^cells, randomized when reaching 50 mm^3^ in groups and treated bi-weekly via intraperitoneal injection with (+BEV, n=6) or without (n=6) BEV (100 µg/dose). Survival end-point: xenografts reaching 500 mm^3^. **(e)** Excised tumor volumes of OV-90^−/−^ xenografts treated with BEV. **(f)** HIF1a western blot analysis in representative OV-90^+/+^ and OV-90^−/−^ xenografts, treated with or without BEV, n=3 per group. Total protein was used as loading control. **(g)** Immunofluorescent staining of endomucin (Endo) and Smooth Muscle Actin (SMA) in OV-90^+/+^ and OV-90^−/−^ xenografts treated with (+BEV) or without BEV. **N**uclei were stained with DAPI, scale bar **5**0 µm. Representative images are shown, together with quantification of total (Endo^+^) and pericyte negative (Endo^+^SMA^−^) vessels.

## Discussion

Our data point to mitochondrial respiration as a vulnerability in quickly growing, aggressive OC cells such as OV-90, which derive from the ascitic fluid of a HGSOC, a stage preluding to pelvic metastatic dissemination^16^. It is interesting that, among solid cancers, metabolism has been used to profile OC into different molecular subsets, accounting for at least one very distinctive aspect of inter-patient heterogeneity. Indeed, OCs have been categorized into high- and low-OXPHOS^17^, high- and low-PGC1 expressing^18^, and glucose deprivation sensitive or resistant^19^, suggesting that there may be specific groups of patients benefiting from a therapeutic effort aimed at inhibiting oxidative phosphorylation through CI. This is relevant in HGSOC since current standard protocols, which envisage the administration of carboplatin and taxanes with or without anti-angiogenic drugs or PARP inhibitors, fail to eradicate the disease in over half the cases, too often leading to chemoresistance onset and fatal recurrence^8,20^. Therefore, the quest for novel adjuvant or neoadjuvant treatment strategies is mandatory and worth pursuing, particularly to prolong the time to relapse through a sustained suppression of cancer cells proliferation. It ought not to discourage that potent CI inhibitors like IACS-010759 fail in trial due to neurotoxicity^21^, or that non-specific metformin yields inconclusive outcomes^22^: we are just beginning to uncover the cell and molecular responses to a spectrum of specific novel CI inhibitors. In this scenario, the use of cell models deprived of CI, while showing this to be a promising approach in OC treatment, will also contribute to depicting the adaptive mechanisms to circumvent in order to gain full efficacy.

Among the vulnerabilities of CI abolishment, one that we robustly and consistently showed to ensue across diverse solid tumor types is the inability to respond to hypoxia^10^. Since hypoxia is the predominant stimulus for neoangiogenesis, driven by a HIF1-dependent VEGF production, we were able to confirm a linear hypothesis of increased sensitivity to anti-VEGF bevacizumab of CI-deprived tumors. These findings corroborate what we previously observed in patient-derived xenografts harboring mitochondrial DNA mutations and impaired oxidative phosphorylation^15^, which our KO models faithfully phenocopy, in terms of a greater sensitivity to bevacizumab. Anti-VEGF therapy is currently administered to subsets of HGSOC patients, mainly in maintenance protocols, with trials both completed and ongoing in combination with PARP, immune checkpoint and, recently, folate receptor inhibitors^23^, to mention a few. The most cogent issue likely concerns both platinum-sensitive and, even more so, platinum-resistant relapsing disease, where only modest, albeit significant, increase in progression-free survival is achieved when BEV is used in combination, in the range of +2 to 4 months^23^. In this dramatic scenario, any low-toxicity therapeutic agent that may boost anti-VEGF approaches, allowing possibly to lower dosage and limit adverse effects, has the potential to make a difference in the fate of relapsing HGSOC patients. As has been seminally suggested in other cancer types^24^, targeting CI compounds may be the class of agents sought for this purpose in HGSOC, for the biological reasons to which our data point.

## Materials and Methods

### Cell line

The human HGSOC cell line OV90^25^ was purchased from ATCC® (Manassas, VA, USA #CRL-3585). Cells were cultivated in Dulbecco’s modified Eagle medium (DMEM) high glucose (Euroclone #ECM0749L), supplemented with 10% FBS (Euroclone #ECS0180L), L-glutamine (2 mM, Euroclone #ECB3000D), penicillin/streptomycin (1×, Euroclone #ECB3001D), uridine (50 μg/mL, Sigma-Aldrich #U3003), in an incubator with a humidified atmosphere at 5% CO2 and 37 °C.

### Genome editing for NDUFS3 knock-out generation

The CRISPR/Cas9 system was used to insert a frameshift mutation in the *NDUFS3* gene in the OV90 cell line. Synthetic RNA guides were designed and purchased from IDT (Alt-R® CRISPR-Cas9 crRNA, IDT). Exon 2 targeting guide TGAACTTGTTGGACATACTT was used. CrRNA and fluorescence-labelled tracrRNA were incubated at 95°C for 5 min and left to cool at room temperature (RT) to form the guide RNA complex. Then, Cas9 protein (Alt-R™ S.p. HiFi Cas9 Nuclease V3, IDT #1081060) was added, together with Cas9 Plus reagent (Invitrogen #CMAX00008) for 5 min to form ribonucleoprotein (RNP) complex. RNP complex was mixed with Lipofectamine CRISPRMAX Cas9 Transfection reagent (Invitrogen #CMAX00008) and placed in a 96-well plate, following the manufacturer’s instructions. Finally, OV90 cells were immediately seeded in the RNP complex containing plate (40K cells/well) and incubated for 48h. Clonal selection was performed by manual cell picking to identify the cells with disruptive *NDFUS3* mutations. Selection media used for the single-cell growth in 96-well plates was composed of DMEM high glucose (Euroclone #ECM0749L), supplemented with 20% FBS (Euroclone #ECS0180L), 2mM L-glutamine (Euroclone #ECB3000D), penicillin/streptomycin (Euroclone #ECB3001D) and uridine (50 μg mL/1, Sigma-Aldrich #U3003). DNA extraction from 96 well plates was performed using 10 μL of Lysis Solution (Sigma-Aldrich #L3289) and 90 μL of Neutralization Buffer (Sigma-Aldrich #N9784) per sample. *NDUFS3* genotype was confirmed by Sanger sequencing using KAPA2G Taq polymerase (Kapa Biosystems #KK5601) and Big Dye protocol (Life Technologies #4337451) using forward (TCTCAAGGTGCTTCAGGGAG) and reverse (GAAACAAGTCTGCCCACTCC) primers with following PCR parameters: 35 cycles of 96°C for 10 sec, 50°C for 5 sec and 60°C for 4 min. After the validation of OV90^−/−^ clones, three of them pooled for the knockout model generation. Likewise, three wildtype counterparts (OV90^+/+^) clones were also pooled to generate control model.

### Generation of cells with inducible NDUFS3 knockout

A stable transgenic cell model that re-expresses *NDUFS3* to allow its inducible knockout was created using the Retro-X Tet-Off©Advanced Inducible Expression system (Clontech #632105). The system is based on gene expression regulation of the *E. coli* tetracycline-resistance operon, regulated by Tet repressor (TetR)^10^. The first element that composes the Tet-Off inducible expression system is a pRetroX-Tet-Off Advanced vector, which encodes the tetracycline-controlled transactivator (tTA), a regulator that activates the transcription of the downstream gene when it binds the promoter tetO sequences. The other element of Tet-Off system is a pRetro-X-Tight-Pur vector, which contains the inducible Tet-responsive PTight promoter upstream of *NDUFS3* gene. The promoter contains a Tet-Responsive-Element (TRE) with seven tetO sequences and located upstream of a modified CMV promoter. In presence of Tet or its derivates, such as doxycycline (DOX) tTA does not bind tetO sequences causing transcriptional repression of the downstream gene. Each Retro-X System vector, along with a viral envelope expression vector, was transiently co-transfected into packaging cells to generate retrovirus stocks, following manufacturer instructions. Target cells were coinfected with the two retrovirus particles and selected simultaneously with previously determined 72-hour killing concentration of G418 (200 μg/ml) and puromycin (0.125 μg/ml), producing the double-stable cell line that expresses the *NDUFS3* gene in the absence of DOX. Single cell cloning of selected cells was then performed, and in the grown colonies NDUFS3 protein expression was analysed by western blot in both total and mitochondrial fractions in presence/absence of 100 ng/mL DOX (Doxycycline Hyclate, =98% (HPLC) D9891-25G). Moreover, to verify the functional effect of inducible NDUFS3 KO on complex I (CI) activity in single clones, they were treated with 100 ng/mL DOX (Doxycycline Hyclate, =98% (HPLC) D9891-25G) for 12-20 days in FBS Tet-free medium (Tet System Approved FBS, US-Sourced 50 mL, Diatech Lab Line, 631105). Subsequently, CI activity was evaluated on mitochondrial fraction with Blue-native polyacrylamide gel electrophoresis (BN-PAGE) followed by CI In-Gel Activity (CI-IGA assay). Clones with both *NDUFS3* expression and functional inducible CI-KO were pulled together creating OV90^−/−S3^ cell model. Similarly, CI-deficient counterpart model was created, by transducing cells with empty vector (OV90^−/−M^). Both cell lines cells were then cultured *in vitro* in Dulbecco’s Modified Eagle Medium (DMEM) containing 4.5 g/L D-Glucose and 0.11 g/L Sodium Pyruvate added with 10% FBS, 2 mM L-Glutamine, 50 μg/mL Uridine, 100 units/mL Penicillin and 100 μg/mL Streptomycin, supplemented with 100 μg/mL G418 and 0.25 μg/mL Puromycin, to sustain the selective pressure.

### SDS-PAGE and Western blot

Whole-cell lysates were obtained by resuspending cell pellets in RIPA lysis buffer containing 50 mM Tris-HCl pH 7.4, 150 mM NaCl, 1% SDS, 1% Triton, 1 mM EDTA pH 7.6, proteinase (Sigma-Aldrich, #04693116001) and phosphatase (Thermo Fisher Scientific, #A32957) inhibitors. The resuspended pellets were incubated for 10 min on ice and frozen/thawed twice. Then, the lysates were centrifuged at 13000 rpm for 10 min at 4°C and the supernatants collected. Protein concentration in the extracts was determined by Lowry protein assay (Bio-Rad #5000116). 50 μg of total protein were combined to 1X Laemmli sample buffer (TrisHCl 63 nM, pH 6.8, glycerol 10%, SDS 2%, β-mercaptoethanol 5%, bromophenol blue 0.0025%), incubated for 5 min at 100° and loaded in 7-10-12% polyacrylamide gels based on the molecular weight of the target (TGX Stain Free TM Fast Cast TM Acrylamide Solution’s kit Biorad.). SDS-PAGE was carried out at 100 V and RT in Tris-Glycine-SDS running buffer (Bio-Rad, #1610772). Precision Plus ProteinTM Standard (Bio-Rad, #1610374) was used for protein molecular weight estimation. Proteins resolved by SDS-PAGE were electroblotted onto 0.45 μm nitrocellulose membranes (Bio-Rad, #1620115) using Trans-Blot® Turbo™ Transfer System (Bio-Rad). Total protein staining was performed in both gel and membrane, and used as loading control when specified. Membranes were rinsed in Tris-Buffered Saline-0.05% Tween 20 (TBS-T) for five min and the non-specific binding sites were saturated by 30 min incubation with 5% powdered skimmed milk in TBS-T at RT. Subsequently, immunodetection of the proteins was performed using commercial protein-specific primary antibodies diluted in 5% milk in TBS-T. Primary antibodies were incubated using the following dilutions/conditions: anti-NDUFS3 (abcam #177471) 1:1000 overnight at 4°C, anti-HIF1α (GTX127309) 1:1000 overnight at 4°C, anti-HSP70 (BD transduction #H53220) 1:1000 x 1h at 4°C, anti-Vinculin (Sigma #V9131) 1:1000 x 1h at RT, and anti-calreticulin (Sigma #C4606) 1:2000 x 1h30’ at RT. Then, washes were performed 4×5 min, using TBS-Tween, and incubation with secondary antibodies (Jackson ImmunoResearch Laboratories #111035144 and #111035146), anti-rabbit diluted 1:10000, anti-mouse diluted 1:20000 in TBS-Tween, was performed for 1h at RT. Next, three washes of ten min in PBS-T were performed. To detect the immunoreactive bands, ClarityTM Western ECL (Bio-Rad, #1705061) was used, and images were developed by exposing in ChemiDocTM MP System (Biorad). Band signal intensities were quantified by densitometry using ImageJ.

### Mitochondria preparation

Mitochondria-enriched fractions were obtained by subcellular fractionation (5–10×10^6^ cells) in presence of digitonin (4 mg/mL x 10 min at 4°C). After removing of digitonin through washing with cold PBS and centrifugation at 4°C at 16,000 × g for 10 min, the samples were resuspended in cold PBS and centrifuged again at 4°C at 16,000 × g for 10 min to precipitate the mitochondria. Crude mitochondria were isolated from snap-frozen xenograft samples. Briefly, at least 30 μg of tissue were disaggregated in sucrose–mannitol buffer (200 mM mannitol, 70 mM sucrose, 1 mM EGTA and 10 mM Tris–HCl at pH 7.6) using Ultra Turrax^®^ homogenizer followed by cells homogenization using a glass/Teflon Potter-Elvehjem homogenizer. Differential centrifugation (600 × g for 10 min at 4°C followed by 10 000 × g for 20 min at 4°C) was performed to separate crude mitochondria from other subcellular fractions. The resulting pellets were stored at −80°C and used for BN-PAGE to analyse NDUFS3 expression and CI-IGA.

Blue-native polyacrylamide gel electrophoresis (BN-PAGE) and CI In-Gel Activity (CI-IGA) Mitochondrial pellets were solubilised resuspending pellet in 100 μl of Mitochondrial Solubilization Buffer (1.5 M aminocaproic acid, 50 mM Bis-Tris/HCl at pH 7) and quantified by Bradford assay. Membrane proteins were solubilised with 2.5 μg DDM/μg total proteins and incubated at 4°C for 5 min. Finally, the samples were centrifuged at 16,000 × g at 4 C° for 30 min to remove the insoluble material, the supernatant was transferred in a new tube, and proteins were quantified by Bradford assay. OXPHOS complexes were separated in their native form by BN-PAGE. Samples containing 30 μg of proteins were prepared in BN-PAGE Sample Buffer 10X (750 mM aminocaproic acid, 50 mM Bis-Tris/HCl pH 7, 0.5 mM EDTA and 5% Coomassie Blue G250) and separated on NuPage 3-12% Bis-Tris gels (Life Technologies) at 150 V and 4 C° for 4h. For separation, Cathode Buffers A (50 mM Tricine, 0.02% Serva Blue-G250, 15 mM Bis-Tris, pH 7), Cathode Buffer B (50 mM Tricine, 0.002% Serva Blue-G250, 15 mM Bis-Tris, pH 7) and Anode Buffer (50 mM Bis-Tris, pH 7) were used. The gel was subsequently used to perform CI-In Gel Activity (CI-IGA) assay or transferred onto a polyvinylidene difluoride (PVDF) membrane for immunodetection of CI. CI-IGA was performed 20 min at RT a BN-PAGE gel with a 10 mL solution containing 2 mg NADH, (Sigma-Aldrich, #N8129), 2.5 mg/mL 3-(4,5-dimethylthiazol-2-yl)-2,5-diphenyltetrazolium bromide (MTT) and 50 μL of Tris/HCl at pH 7.4. PVDF membrane was blocked 1 hour at RT with 5% powdered skimmed milk in PBS containing 0.1% Tween 20 (PBS-T) and incubated with the following primary antibodies: anti-NDUFS3 (Abcam #177471) 1:1000 overnight at 4°C; anti-HSP60 (Santa Cruz Biotechnology, #sc-13966) 1:1000 for 2 hours at RT. Secondary antibody (Jackson ImmunoResearch Laboratories, #111035144) was incubated for 40 min at RT using 1:5000 dilution in PBS-T. Membrane was developed using Clarity Western ECL Substrate (Bio–Rad #1705061), and ECL was detected with ChemiDoc (Bio–Rad).

### Xenograft growth

In vivo studies were performed at Department of Medical and Surgical Sciences, at the CRBA animal facility of the University of Bologna, and at Department of Surgery, Oncology and Gastroenterology, University of Padua. All relevant ethical regulations for animal testing and research have been complied. Procedures involving animals and their care were conformed to institutional guidelines that comply with national and international laws and policies (EEC Council Directive 2010/63/EU, OJ L 276, 20.10.2010) and were authorized by the Italian Ministry of Health (Authorization n. 1109/2024-PR and n. 77/2022-PR). NOD/SCID mice were purchased from Charles River Laboratories. Five to six-week-old female mice were subcutaneously injected with a 100 μL suspension of 1.5×10^6^ cells in serum free medium and matrigel (Corning #356234) in the right flank of the animal. Xenograft size was measured with a sliding caliper twice a week and calculated according to the formula: volume = width × height × length/2. Mice were sacrificed either simultaneously, when the first xenograft reached 10% of animal weight (Figure 1) or consecutively, when each animal reached xenograft volume corresponding >500 mm^3^ of animal weight (Figure 2, Figure 4). For the DOX-induced experiment, mice were sequentially randomized in DOX-treated and control group when tumor was palpable (50 mm^3^), and 3% sucrose with or without DOX (1 mg/mL) was added into the drinking water. The animals were similarly randomized for the experiment involving bevacizumab and treated bi-weekly intraperitoneally with either PBS or anti-human VEGF mAb (bevacizumab-OYAVAS) at 100 µg/dose.

### Immunohistochemical staining

The samples were formalin fixed and paraffin embedded following standard protocols. Tissue sections (4 μm) were deparaffinized in toluene and rehydrated by a series of 2-3 min washes in 100%, 96%, and 70% ethanol and distilled water followed by heat-induced epitope retrieval in TE-T buffer (10 mM Tris pH 8.0, 1 mM EDTA, 0.05% Tween 20) for 1 h at 95 °C and 20 min at RT, using controlled temperature pressure cooker, and then sections were equilibrated with water. Then, sections were equilibrated with phosphate-buffered saline containing 0.5% Tween 20 (PBS-T pH 7.4). Primary antibodies were diluted in Antibody diluent with background reducing components (Dako #S3022) and incubated at RT for 1 h. The following primary antibodies were used: rabbit monoclonal anti-NDUFS3 (1:200, Abcam #177471); mouse monoclonal anti-NDUFS4 (1:1000, Abcam #55540) and mouse monoclonal anti-KI-67 (1:100, Dako #M7240). Blocking, secondary antibody staining, and development were performed using the Envision Detection System (Dako #K4065). To visualise the antigen, 3,3-diaminobenzidine tetrahydrochloride (DAB) (20 μl chromogen x 1 mL substrate buffer; Dako Diagnostic) was added at RT. Slides were counterstained with haematoxylin for 5 min, rinsed in tap water, dehydrated, placed in toluene, and mounted. Alternatively, automatic immunostainer BOND RX processing module (Leica Biosystems #3219740) was used with Bond Polymer Refine Detection kit (Leica Biosystems #DS9800) and following protocol: Marker 30 min, Peroxide Block 15 min, Post Primary 10 min, Polymer 10 min, Mixed DAB Refine 10 min, Hematoxylin 7 min. Images were obtained by using the Leica DM750 microscope equipped with the Leica ICC50 W camera and with SLIDEVIEW VS200. Alternatively, Olympus SLIDEVIEW VS200 slide scanner and Olympus image viewer OlyVIA were used for image acquisition and analysis. For evaluation of KI-67 positive nuclei, cells were counted by two independent operators at 60x magnification in 8-10 fields of view (FOV) per tumor. For NDUFS3 and NDUFS4 positivity was evaluated by two independent operators at 10X magnification in 5-6 FOV.

### Masson’s Trichrome coloration

For collagen fibers visualization and distribution in tissue, Masson’s Trichrome Staining Kit (Polysciences #25088) was used according to manufacturer instructions. Briefly, after sections dewaxing and hydration, slides were incubated for 1h at 60°C in Bouin’s fixative (#25088A) and then washed in tap water for 5 min. Slides were stained for 10 min in Weigert’s working haematoxylin solution, prepared by mixing Weigert’s Hematoxylin A (#25088B1) and Weigert’s Hematoxylin B (#25088B2) at a 1:1 ratio, washed in tap water for 5 min, and then stained with Biebrich Scarlet - Acid Fuchsin Solution (#25088C) for 5 min. After a rinse, slides were incubated in Phosphomolybdic Acid (#25088D) for 10 min and transferred directly into Alanine Blue (#25088E) for 5 min, following a rinse in water. Lastly, slides were transferred to 1% Acetic Acid (#25088F) for 1 min, rinsed in water and dehydrated by a series of 2-3 min whases in 70%, 96%, 100% ethanol and toluene, then mounted. Images were obtained by using the Leica DM750 microscope equipped with the Leica ICC50 W camera. For the analysis, stromal percentages in 3 FOV per sample were quantified at 4X magnification by two independent operators.

### Quantitative Real-Time PCR

Total RNA was extracted from snap frozen xenograft samples using the RNeasy Mini Kit (QIAGEN, Cat. No. 74106) according to the manufacturer’s instructions. RNA was quantified using NanoDrop™ 2000 Spectrophotometer (Thermo Scientific). For gene expression analysis, 200 ng of total RNA were reverse-transcribed into cDNA using the High-Capacity cDNA Reverse Transcription Kit (Applied Biosystems, Cat. No. 4368814), following the manufacturer’s protocol. Quantitative real-time PCR (qRT-PCR) was performed to evaluate the expression of the following HIF1 target genes: *GLUT1* (forward: ACTCCATCATGGGAACAAG; reverse: TCTGCCGACTCTCTTCCTTC), *VEGFA* (forward: ACGAGGGCCTGGAGTGTGT; reverse: CGCATAATCTGATGGTGATG), *LDHA* (forward: TGGGAGTTCACCCATTAAGC reverse: AGCACTCTCAACCACCTGCT), and *MIF* (forward: AGAACCGCTCCTACAGCAAG; reverse: GAGTTGTTCCAGCCCACATT). Reactions were carried out using SYBR® Green chemistry (Promega) with GoTaq® qPCR Master Mix (Promega, Cat. No. A6002) on a 7500 Fast Real-Time PCR System (Applied Biosystems). The thermal cycling conditions were as follows: 95 °C for 5 min, followed by 45 cycles of 95 °C for 15 s and 60 °C for 45 s. Relative gene expression was calculated using the 2^-ΔΔCt method. ΔCt values were obtained by normalizing the Ct of each target gene to the reference gene TBP [ΔCt = Ct(target) - Ct(TBP)]. ΔΔCt values were calculated as ΔCt(experimental) - ΔCt(control), where the control corresponded to the mean ΔCt of the control group.

### Ultrasound examination and image analysis

US imaging of the xenograft masses was done using the Esaote MYLAB 70 XVG US machine (Esaote, Genova, Italy). All animals were sedated with a continuous flow of isoflurane connected to a charcoal gas scavenging system and placed in prone position on a heated mat to ensure the temperature homeostasis was maintenance throughout the procedure. The hair covering the tumor was shaved with an electric razor and a generous layer of preheated ultrasound gel was applied so to avoid significant external pressure on the tumor that could alter blood flow. First, B-mode US with a linear probe LA523, REF 9600156000 (4.0 - 13.0 mHz) was used to define the appearance of the xenograft after which power Doppler was used to search for tumoral blood flow. Multiple planes and angles were scanned to ensure an optimal signal.

ImageJ^26^ was used to assess and quantify the intratumoral Doppler signal, the pixels corresponding to the colored area – red or blue - within the tumor were selected and measured to obtain an indirect estimation of the blood flow area. The resulting value was represented as a percentage of the total tumor area measured on the same US image.

### Immunofluorescence

The xenografts were formalin-fixed following standard protocols. Tissue sections (4 μm) were deparaffinised in toluene and rehydrated by a series of 2-3 min washes in 100%, 96%, and 70% ethanol and distilled water. The rehydration was followed by citrate antigen retrieval (10 mM sodium citrate, pH 6) for 10 min at 95 °C, 15 min at 60 °C and 20 min at RT, for vessels analysis, or trypsin-based antigen retrieval (0.05% trypsin in 0.1 mM Calcium chloride solution, pH 8.2) performed for 30 min at 37 °C for macrophage analysis. Blocking was performed with goat serum 10% (Abcam #156046) for 10 min at RT. Rat anti-Endomucin (1:200, Santa Cruz #SC-65495), mouse anti-SMA (1:750, Dako #M0851) and Anti-mouse F4/80 (1:100, eBiosciences #14-4801) primary antibodies were used. Slides were washed in PBS and incubated with Alexa Fluor secondary antibodies 488 Goat anti Mouse diluted 1:500 (Thermo Fisher Scientific #A-11017) and 555 Goat anti Rat diluted 1:350 (Thermo Fisher Scientific #A-21434) for 40 min at RT in the dark. Slides were mounted with ProLongTM gold antifade reagent with DAPI (Thermofisher #1804483). Images were taken with SLIDEVIEW™ VS200 slide scanner (Olympus Corporation, Tokyo, Japan). Macrophages (F4/80^+^) were counted at 20X magnification in 3-5 fields of view per tumor by two independent operators. Mature vessels (Endo^+^SMA^+^) and immature vessels (Endo^+^SMA^−^) were counted in 3-5 fields of view at ×20 magnification

### Statistical analysis

GraphPad Prism version 8 (GraphPad Software Inc., San Diego, CA, USA) was used to perform statistical tests and create bar plots and graphs. Grubbs test was applied to identify outliers. If found, the outliers were indicated in the graph but excluded from the statistical comparison. Unless stated otherwise, two-tailed unpaired Student’s t-tests assuming equal variance were performed to compare means. Shapiro-Wilk test was applied to test the normality of data distribution. If not normally distributed, the data between the groups were compared by using Mann-Whitney test. F-test was used to compare variances between the groups, and when significant, the data were transformed (y′ = logy) prior to the t-test calculus. If the variance difference persisted despite the transformation, Welch’s correction was applied. Survival curves were estimated using Kaplan–Meier product-limit method and compared using a log-rank test (Mantel–Cox).

## Acknowledgements

We are grateful to drs. Monica De Luise, Silvia Lemma, Maurizio Baldassarre and Sara Coluccelli for technical assistance.

## Author contributions

Conceptualization, I.K., A.M.P., G.G.; methodology, I.K., B.C., F.N., S.C, M.S., C.A.C., M.G., L.S., S.M., E.L., E.A., L.I.; acquisition and analysis, I.K., B.C., F.N., S.C, C.A.C., M.G., L.S., S.M., E.L., E.A., L.I.; data interpretation: I.K., B.C., M.S., L.I., A.M.P., G.G.,; writing – original draft, I.K., G.G.; writing – review & editing, I.K., B.C., F.N., S.C, M.S., C.A.C., S.M., L.I., A.M.P., S.I., G.G; project administration, G.G, funding acquisition and resources, I.K., A.M.P., S.I., G.G. All authors have approved the submitted version of the work and have agreed both to be personally accountable for the author’s own contributions and to ensure that questions related to the accuracy or integrity of any part of the work, even ones in which the author was not personally involved, are appropriately investigated, resolved, and the resolution documented in the literature.

## References

1. Leone, G., Abla, H., Gasparre, G., Porcelli, A. M. & Iommarini, L. The Oncojanus Paradigm of Respiratory Complex I. Genes 9, 243 (2018).

2. Birsoy, K., Wang, T., Chen, W. W., Freinkman, E., Abu-Remaileh, M. & Sabatini, D. M. An Essential Role of the Mitochondrial Electron Transport Chain in Cell Proliferation Is to Enable Aspartate Synthesis. Cell 162, 540–551 (2015).

3. Porcelli, A. M., Ghelli, A., Ceccarelli, C., Lang, M., Cenacchi, G., Capristo, M., Pennisi, L. F., Morra, I., Ciccarelli, E., Melcarne, A., Bartoletti-Stella, A., Salfi, N., Tallini, G., Martinuzzi, A., Carelli, V., Attimonelli, M., Rugolo, M., Romeo, G. & Gasparre, G. The genetic and metabolic signature of oncocytic transformation implicates HIF1alpha destabilization. Hum. Mol. Genet. 19, 1019–1032 (2010).

4. Molina, J. R., Sun, Y., Protopopova, M., Gera, S., Bandi, M., Bristow, C., McAfoos, T., Morlacchi, P., Ackroyd, J., Agip, A.-N. A., Al-Atrash, G., Asara, J., Bardenhagen, J., Carrillo, C. C., Carroll, C., Chang, E., Ciurea, S., Cross, J. B., Czako, B., Deem, A., Daver, N., de Groot, J. F., Dong, J.-W., Feng, N., Gao, G., Gay, J., Do, M. G., Greer, J., Giuliani, V., Han, J., Han, L., Henry, V. K., Hirst, J., Huang, S., Jiang, Y., Kang, Z., Khor, T., Konoplev, S., Lin, Y.-H., Liu, G., Lodi, A., Lofton, T., Ma, H., Mahendra, M., Matre, P., Mullinax, R., Peoples, M., Petrocchi, A., Rodriguez-Canale, J., Serreli, R., Shi, T., Smith, M., Tabe, Y., Theroff, J., Tiziani, S., Xu, Q., Zhang, Q., Muller, F., DePinho, R. A., Toniatti, C., Draetta, G. F., Heffernan, T. P., Konopleva, M., Jones, P., Di Francesco, M. E. & Marszalek, J. R. An inhibitor of oxidative phosphorylation exploits cancer vulnerability. Nat. Med. 24, 1036–1046 (2018).

5. Wheaton, W. W., Weinberg, S. E., Hamanaka, R. B., Soberanes, S., Sullivan, L. B., Anso, E., Glasauer, A., Dufour, E., Mutlu, G. M., Budigner, G. S. & Chandel, N. S. Metformin inhibits mitochondrial complex I of cancer cells to reduce tumorigenesis. eLife 3, e02242 (2014).

6. Urpilainen, E., Puistola, U., Boussios, S. & Karihtala, P. Metformin and ovarian cancer: the evidence. Ann. Transl. Med. 8, 1711 (2020).

7. Zhang, X., Shetty, M., Clemente, V., Linder, S. & Bazzaro, M. Targeting Mitochondrial Metabolism in Clear Cell Carcinoma of the Ovaries. Int. J. Mol. Sci. 22, 4750 (2021).

8. Mahmood, R. D., Morgan, R. D., Edmondson, R. J., Clamp, A. R. & Jayson, G. C. First-Line Management of Advanced High-Grade Serous Ovarian Cancer. Curr. Oncol. Rep. 22, 64 (2020).

9. Li, T., Zhang, H., Lian, M., He, Q., Lv, M., Zhai, L., Zhou, J., Wu, K. & Yi, M. Global status and attributable risk factors of breast, cervical, ovarian, and uterine cancers from 1990 to 2021. J. Hematol. Oncol.J Hematol Oncol 18, 5 (2025).

10. Kurelac, I., Iommarini, L., Vatrinet, R., Amato, L. B., De Luise, M., Leone, G., Girolimetti, G., Umesh Ganesh, N., Bridgeman, V. L., Ombrato, L., Columbaro, M., Ragazzi, M., Gibellini, L., Sollazzo, M., Feichtinger, R. G., Vidali, S., Baldassarre, M., Foriel, S., Vidone, M., Cossarizza, A., Grifoni, D., Kofler, B., Malanchi, I., Porcelli, A. M. & Gasparre, G. Inducing cancer indolence by targeting mitochondrial Complex I is potentiated by blocking macrophage-mediated adaptive responses. Nat. Commun. 10, 903 (2019).

11. D’Angelo, L., Astro, E., De Luise, M., Kurelac, I., Umesh-Ganesh, N., Ding, S., Fearnley, I. M., Gasparre, G., Zeviani, M., Porcelli, A. M., Fernandez-Vizarra, E. & Iommarini, L. NDUFS3 depletion permits complex I maturation and reveals TMEM126A/OPA7 as an assembly factor binding the ND4-module intermediate. Cell Rep. 35, 109002 (2021).

12. De Luise, M., Sollazzo, M., Lama, E., Coadă, C. A., Bressi, L., Iorio, M., Cavina, B., D’Angelo, L., Milioni, S., Marchio, L., Miglietta, S., Coluccelli, S., Tedesco, G., Ghelli, A., Lemma, S., Perrone, A. M., Kurelac, I., Iommarini, L., Porcelli, A. M. & Gasparre, G. Inducing respiratory complex I impairment elicits an increase in PGC1α in ovarian cancer. Sci. Rep. 12, 8020 (2022).

13. Zimmermann, F. A., Mayr, J. A., Feichtinger, R., Neureiter, D., Lechner, R., Koegler, C., Ratschek, M., Rusmir, H., Sargsyan, K., Sperl, W. & Kofler, B. Respiratory chain complex I is a mitochondrial tumor suppressor of oncocytic tumors. Front. Biosci. 3, 315–325 (2011).

14. Mateiou, C., Lokhande, L., Diep, L. H., Knulst, M., Carlsson, E., Ek, S., Sundfeldt, K. & Gerdtsson, A. Spatial tumor immune microenvironment phenotypes in ovarian cancer. Npj Precis. Oncol. 8, 148 (2024).

15. Boso, D., Piga, I., Trento, C., Minuzzo, S., Angi, E., Iommarini, L., Lazzarini, E., Caporali, L., Fiorini, C., D’Angelo, L., De Luise, M., Kurelac, I., Fassan, M., Porcelli, A. M., Navaglia, F., Billato, I., Esposito, G., Gasparre, G., Romualdi, C. & Indraccolo, S. Pathogenic mitochondrial DNA variants are associated with response to anti-VEGF therapy in ovarian cancer PDX models. J. Exp. Clin. Cancer Res. CR 43, 325 (2024).

16. Rickard, B. P., Conrad, C., Sorrin, A. J., Ruhi, M. K., Reader, J. C., Huang, S. A., Franco, W., Scarcelli, G., Polacheck, W. J., Roque, D. M., Del Carmen, M. G., Huang, H.-C., Demirci, U. & Rizvi, I. Malignant Ascites in Ovarian Cancer: Cellular, Acellular, and Biophysical Determinants of Molecular Characteristics and Therapy Response. Cancers 13, 4318 (2021).

17. Gentric, G., Kieffer, Y., Mieulet, V., Goundiam, O., Bonneau, C., Nemati, F., Hurbain, I., Raposo, G., Popova, T., Stern, M.-H., Lallemand-Breitenbach, V., Müller, S., Cañeque, T., Rodriguez, R., Vincent-Salomon, A., de Thé, H., Rossignol, R. & Mechta-Grigoriou, F. PML-Regulated Mitochondrial Metabolism Enhances Chemosensitivity in Human Ovarian Cancers. Cell Metab. 29, 156–173.e10 (2019).

18. Ghilardi, C., Moreira-Barbosa, C., Brunelli, L., Ostano, P., Panini, N., Lupi, M., Anastasia, A., Fiordaliso, F., Salio, M., Formenti, L., Russo, M., Arrigoni, E., Chiaradonna, F., Chiorino, G., Draetta, G., Marszalek, J. R., Vellano, C. P., Pastorelli, R., Bani, M., Decio, A. & Giavazzi, R. PGC1α/β Expression Predicts Therapeutic Response to Oxidative Phosphorylation Inhibition in Ovarian Cancer. Cancer Res. 82, 1423–1434 (2022).

19. Boso, D., Tognon, M., Curtarello, M., Minuzzo, S., Piga, I., Brillo, V., Lazzarini, E., Carlet, J., Marra, L., Trento, C., Rasola, A., Masgras, I., Caporali, L., Del Ben, F., Brisotto, G., Turetta, M., Pastorelli, R., Brunelli, L., Navaglia, F., Esposito, G., Grassi, A. & Indraccolo, S. Anti-VEGF therapy selects for clones resistant to glucose starvation in ovarian cancer xenografts. J. Exp. Clin. Cancer Res. CR 42, 196 (2023).

20. Polajžer, S. & Černe, K. Precision Medicine in High-Grade Serous Ovarian Cancer: Targeted Therapies and the Challenge of Chemoresistance. Int. J. Mol. Sci. 26, 2545 (2025).

21. Yap, T. A., Daver, N., Mahendra, M., Zhang, J., Kamiya-Matsuoka, C., Meric-Bernstam, F., Kantarjian, H. M., Ravandi, F., Collins, M. E., Francesco, M. E. D., Dumbrava, E. E., Fu, S., Gao, S., Gay, J. P., Gera, S., Han, J., Hong, D. S., Jabbour, E. J., Ju, Z., Karp, D. D., Lodi, A., Molina, J. R., Baran, N., Naing, A., Ohanian, M., Pant, S., Pemmaraju, N., Bose, P., Piha-Paul, S. A., Rodon, J., Salguero, C., Sasaki, K., Singh, A. K., Subbiah, V., Tsimberidou, A. M., Xu, Q. A., Yilmaz, M., Zhang, Q., Li, Y., Bristow, C. A., Bhattacharjee, M. B., Tiziani, S., Heffernan, T. P., Vellano, C. P., Jones, P., Heijnen, C. J., Kavelaars, A., Marszalek, J. R. & Konopleva, M. Complex I inhibitor of oxidative phosphorylation in advanced solid tumors and acute myeloid leukemia: phase I trials. Nat. Med. 29, 115–126 (2023).

22. He, L. & Wondisford, F. E. Metformin action: concentrations matter. Cell Metab. 21, 159–162 (2015).

23. Lamia, M. R., Perri, E., Baldassarre, G., Pignata, S., D’Alessio, C., Limongello, D. & Basso-Valentina, F. Bevacizumab in ovarian cancer: Clinical data and predictive and prognostic biomarkers. Clin. Transl. Med. 16, e70591 (2026).

24. Marciano, R., Prasad, M., Ievy, T., Tzadok, S., Leprivier, G., Elkabets, M. & Rotblat, B. High-Throughput Screening Identified Compounds Sensitizing Tumor Cells to Glucose Starvation in Culture and VEGF Inhibitors In Vivo. Cancers 11, 156 (2019).

25. Bourgeois, D. L., Kabarowski, K. A., Porubsky, V. L. & Kreeger, P. K. High-grade serous ovarian cancer cell lines exhibit heterogeneous responses to growth factor stimulation. Cancer Cell Int. 15, 112 (2015).

26. Schneider, C. A., Rasband, W. S. & Eliceiri, K. W. NIH Image to ImageJ: 25 years of image analysis. Nat. Methods 9, 671–675 (2012).

